# Pathogenicity and transmissibility of Monkeypox virus in dormice

**DOI:** 10.1101/2025.06.17.660157

**Authors:** Zhaoliang Chen, Lei Zhang, Linzhi Li, Mingjie Shao, Zongzheng Zhao, Chao Shang, Zirui Liu, Juxiang Liu, Yan Liu, Xiao Li, Zhendong Guo

**Author notes:** Authors to whom correspondence should be addressed; E-Mails (Z.G.), (X.L.), (Y.L.); (Z.G.), (X.L.), (Y.L.).

## Abstract

The global spread of the monkeypox virus (MPXV) poses a significant challenge to public health, yet reliable and consistent animal models for evaluating MPXV remain limited, which to some extent restricts the advancement of treatment strategies and transmission-blocking technologies. As natural hosts, African dormice (*Graphiurus spp.*) represent promising candidates. However, the biological characteristics of MPXV in dormice remain largely unexplored. This study systematically evaluated the pathogenicity and transmissibility of MPXV in dormice. Experimental results demonstrated that dormice are highly susceptible to MPXV infection. Following intranasal inoculation, MPXV induced significant weight loss, lethal infections, and multi-organ pathological damage, with robust viral replication in respiratory and liver tissues. Notably, MPXV can efficiently transmit among dormice through direct contact, with one-third of the contact-exposed dormice shedding infectious viruses and two-thirds exhibiting seropositivity. In addition, one-third of the airborne exposure dormice show seropositivity, yet no infectious viruses were detected in their respiratory tissues, indicating that the airborne transmissibility of MPXV among dormice is relatively restricted. Furthermore, it was observed that infected dormice continuously released large quantities of virus-laden aerosols, with emissions peaking on 12 dpi (3.78±1.01×10^6^ copies/dormouse/hour of respiration). Particle size analysis revealed that the viral copies in ≥7 μm coarse particles accounted for >90.9% of exhaled viral aerosols during peak shedding (8–14 dpi). These findings demonstrate that the African dormice serve as an effective animal model for assessing MPXV infection, pathogenicity, and transmission dynamics. Simultaneously, enhanced monitoring of wild dormouse populations is critical due to their potential role in MPXV transmission chains.

## Introduction

The monkeypox virus (MPXV) is a member of the Orthopoxvirus genus within the Poxviridae family and shares the same genus as the variola virus. Additionally, it is classified as a zoonotic pathogen(1, 2).This virus is capable of causing human infection with monkeypox, and its clinical manifestations closely resemble those of smallpox(3). The incubation period for MPXV infection varies among individuals, followed by a prodromal phase lasting 1 to 5 days. Symptoms during this phase may include fever, chills, diaphoresis, headache, fatigue, back pain, malaise, as well as swelling and tenderness of lymph nodes in the cervical, axillary, and/or inguinal regions. A characteristic clinical feature is the development of skin lesions that are similar to those observed in smallpox infections (4, 5). Although the mortality rate among adults is generally low, it can surpass 10% in children and individuals with compromised immune systems(6).

The first documented case of monkeypox can be traced back to 1958, involving a group of crab-eating macaques and rhesus macaques that were primarily imported from Singapore to Copenhagen, Denmark for laboratory research purposes. These primates were subsequently infected with the virus (7). In 1970, Zaire (now the Democratic Republic of the Congo) documented the first confirmed human case of monkeypox globally. Since then, the pathogen has progressively spread across Africa and to other regions, with endemic prevalence primarily observed in Central and West African countries. This epidemiological pattern persisted until 2003 (8, 9). In May 2003, the first cases of monkeypox were reported in several states in the Midwestern United States (10). Subsequently, there has been a yearly increase in the number of cases reported in countries and regions outside the traditional endemic areas. In recent years, the World Health Organization (WHO) has declared monkeypox a “Public Health Emergency of International Concern” on two occasions: in July 2022 and August 2024 (11). Given the severity of the epidemic situation, it is imperative to analyze the transmission routes and patterns of the MPXV so as to facilitate the development of scientifically rigorous and effective prevention and control strategies.

Establishing appropriate animal models represents a critical scientific foundation for elucidating the transmission routes and epidemic patterns of the MPXV. Existing studies have demonstrated that various susceptible animals can serve as viable biological models for investigating the MPXV, enabling the evaluation of its pathogenic mechanisms, vaccine efficacy, and treatment strategies (12). Specifically, mammalian models encompass the African dormouse (13), prairie dog (14), Gambian pouched rat (15), African rope squirrel (16, 17), ground squirrel (18), and rabbit (19, 20). Non-human primate models include Macaca mulatta (21), Macaca fascicularis (22), and marmosets (23). Although inbred mouse strains, such as BALB/c and C57BL/6, exhibit relatively low susceptibility to monkeypox virus infection, their distinct innate immune response characteristics and adaptive immune regulatory mechanisms offer a valuable experimental model for elucidating the pathogenic mechanisms of the monkeypox virus (24). Limited research evidence suggests that the MPXV may spread within black-tailed prairie dog (*Cynomys ludovicianus*) populations through various transmission mechanisms, including direct contact with infected individuals, indirect contact with contaminated materials (e.g., bedding), and respiratory transmission routes (25, 26). However, despite the successful establishment of various animal models for MPXV, apart from the black-tailed prairie dog model, there is limited literature on the transmissibility of the MPXV in other experimental or susceptible animal models. Additional data are required to further investigate the transmissibility of the MPXV.

The African dormouse (*Graphiurus spp.*) is a small rodent that has been identified as highly susceptible to the MPXV, which is capable of replicating efficiently in multiple organs within its body(27). The dormouse not only serves as a natural host for the MPXV but also represents a promising candidate for the development of animal models in research. However, the biological characteristics of the MPXV in African dormice remain largely unexplored, and there is notable lack of studies investigating the transmission dynamics and patterns of the virus within these populations.

In this study, we conducted a comprehensive evaluation of the pathogenicity and transmissibility of the MPXV in African dormice. We first developed a dormouse model of MPXV infection to systematically characterize the pathogenic profile of the virus in this rodent species, including clinical manifestations, virological dynamics, and histopathological changes. Transmissibility assessments were subsequently performed via direct contact and aerosol transmission routes to quantify transmission efficiency and delineate the epidemiological potential of these routes. Additionally, we conducted time-series analyses to elucidate the kinetics of viral aerosol shedding from infected dormice, providing critical insights into environmental virus dissemination. This study demonstrates that African dormice can serve as a reliable animal model for MPXV research, and enhances our understanding of MPXV epidemiology and provides essential evidence to inform public health strategies aimed at mitigating zoonotic spillover and spread.

## Materials and methods

### Ethics statement

All the animal studies were performed in strict accordance with the guidelines for animal welfare of the World Organization for Animal Health. The experimental protocols involving animals were approved by the Animal Care and Use Committee of the Changchun Veterinary Research Institute, Chinese Academy of Agricultural Sciences (approval number: IACUC of AMMS-11-2023-040). All experiments involving the monkeypox virus were conducted in an Animal Biosafety level 3 (ABSL-3) laboratory at the Changchun Veterinary Research Institute.

### Virus and cell

The hMPXV/China/CCVRI-01/2023 strain of the monkeypox virus was generously provided by the Eighth Affiliated Hospital of Guangzhou Medical University and subsequently propagated in Vero E6 cells. This strain was isolated from a patient in Guangzhou, China. The master seed virus, obtained through proliferation, is stored at −80 °C. Vero E6 cells were cultured in a humidified incubator set at 37 °C with 5% CO_2_. The culture medium was composed of Dulbecco’s Modified Eagle Medium (DMEM, Sigma-Aldrich) supplemented with 10% fetal bovine serum (FBS, Invitrogen), 50 U/mL penicillin, and 50 μg/mL streptomycin.

### Animals and viral challenge

The experimental animals utilized in this study were male Spanish dormice (Dryomys nitedula, Rodentia, Myoxidae, Dryomys), aged between 10 and 12 months, with a body weight ranging from 20 to 30 grams. They are reared and cultivated under controlled conditions in the laboratory of the Changchun Veterinary Research Institute. The animals were maintained under strictly controlled environmental conditions, with the temperature set at 24 ± 2°C, relative humidity ranging from 40% to 70%, and a light/dark cycle of 12 hours each. Additionally, food and water were freely available. These animals were randomly allocated into various experimental groups and acclimatized to the ABSL-3 facility for 2–3 days before the experiment commenced. For the viral challenge experimental, animals were anesthetized via isoflurane inhalation. After confirming successful administration of anesthesia, the animals were intranasally inoculated with monkeypox virus at a dose of 50 μL of viral solution containing 10^5.3^ TCID_50_ of the virus.

### Pathogenicity of monkeypox virus in dormice

To monitor the body weight changes and survival rate of dormice following MPXV infection, a total of 20 dormice were randomly allocated into two groups: the infected group and the control group (n=10). The infected group was intranasally inoculated with 10^5.3^ TCID_50_ of MPXV, while the control group was intranasally inoculated with an equal volume of DMEM. The body weight and survival of these dormice were monitored daily for 15 days post-challenge, and dormice that lost more than 25% of their initial body weight were humanely euthanized.

To determine the tissue distribution of MPXV and assess pathological damage in dormice following MPXV infection, twenty-four dormice were assigned to the infected group following intranasal inoculation with MPXV, while three dormice were allocated to the control group after intranasal administration of DMEM. On 3, 5, 7, 9, 11, 13, and 15 days post-infection (dpi), three dormice were randomly selected from the infected group, and their tissues, including nasal turbinate, brain, heart, lung, liver, trachea, kidney, spleen, intestine, and testis, were collected. These tissues were subsequently placed in 1 mL of PBS supplemented with 2% penicillin-streptomycin. Following tissue homogenization, the samples were centrifuged, and the resulting supernatant was carefully collected. The supernatant was subsequently diluted and inoculated into Vero-E6 cells, and the TCID_50_ was determined using the Reed-Muench method. On 7 dpi, tissues were collected from three dormice in the infected group and three in the control group, followed by fixation in paraformaldehyde. Thereafter, the specimens were embedded in paraffin, sectioned, and stained with hematoxylin and eosin (H&E) for histopathological examination.

### Transmissibility of monkeypox virus in dormice

To evaluate the transmissibility of MPXV between dormice via direct contact and airborne transmission, 48 donor dormice were intranasally inoculated with MPXV. At 24 hpi, 24 donor dormice were transferred to four clean cages and co-housed with 24 naïve recipient dormice for direct contact transmission studies. The animals were randomly assigned to four cages, with six donor dormice housed together with six naive direct-contact recipient dormice in each cage. Additionally, the remaining 24 donor dormice were transferred to four wire-frame cages adjacent to 24 naïve recipient dormice for airborne transmission studies. The animals were also randomly assigned to four cages, with six donor dormice housed adjacent to six naïve airborne-exposure recipient dormice in each cage. The distance between the infected dormice and the airborne-exposed dormice was 2 cm, which prevented direct contact while allowing indirect interaction via respiratory transmission.

On 7, 10, 13, and 16 days post-infection or exposure, the nasal turbinate, trachea, and lungs were collected from six donor dormice or six recipient dormice in each group and subsequently homogenized in 1 mL of PBS supplemented with 2% penicillin-streptomycin. Thereafter, the TCID_50_ was determined, and the viral nucleic acid was quantified using a standardized method. On 16 days post-infection or exposure, whole blood was collected from the dormice in each group and incubated overnight at 4 °C to facilitate serum separation. The resulting serum samples were serially diluted and subsequently mixed in a 1:1 ratio with a viral inoculum containing 100 TCID_50_. The mixtures were then incubated in 96-well plates for 1 hour at 37°C, prior to the addition of Vero-E6 cells. Cells were cultured under conditions promoting adherence and growth at 37 °C. Four days later, the neutralizing antibody titer in the serum was determined by evaluating the number of wells exhibiting cytopathic effects (CPE) and the number of wells demonstrating complete antibody-mediated protection against CPE across each serum dilution.

### Collection of exhaled viral aerosols from dormice

Four dormice were intranasally inoculated with MPXV. Exhaled breath samples from the dormice were collected on 2, 4, 6, 8, 10, 12, 14, and 16 dpi using six-stage Anderson impactors (TE-20-800; Tisch Inc., Cleves, OH, USA). The collection parameters were set at a flow rate of 28.3 liters per minute for a duration of 60 minutes per session. Six-stage Anderson impactors can classify aerosol particles in the collected oral and nasal exhaled breath into six size ranges (> 7 μm, 4.7-7 μm, 3.3-4.7 μm, 2.1-3.3 μm, 1.1-2.1 μm, and 0.65-1.1 μm) based on their aerodynamic diameters. Aerosol particles within each size range are subsequently collected onto separate, pre-sterilized gelatin filters (Sartorius, Germany), as previously described(28). The aerosol samples were collected and subsequently divided into two equal portions: one portion was used for the quantitative detection of viral nucleic acid, while the other was employed for viral titer determination using the TCID_50_ method.

### Determination of the median infective dose (ID_50_)

Twenty dormice were randomly allocated into four groups, with five animals per group. The four groups were intranasally inoculated with monkeypox virus at doses of 10^4.3^ TCID_50_, 10^3.3^ TCID_50_, 10^2.3^ TCID_50_, and 10^1.3^ TCID_50_, respectively. On day 7 post-infection, these dormice were euthanized, and their respiratory tissues, including the nasal turbinate, trachea, and lungs, were harvested for viral titer determination. The ID_50_ for instilled MPXV was determined by probit analysis, based on the infection efficiency of dormice in each group following exposure to the respective viral doses.

### Viral nucleic acid detection

DNA was extracted using the TIANamp Virus DNA/RNA Kit (TIANGEN, China) and quantified by quantitative real-time PCR (Q-RT-PCR) using the One Step PrimeScript^TM^ RT-PCR Kit (TaKaRa, Japan), following the manufacturers’ protocols. The primers and probe targeting the F3L gene of monkeypox virus were as follows: [Forward: 5’-CTCATTGATTTTTCGCGGGATA-3’; Reverse: 5’-GACGATACTCCTCCTCGTTGGT-3’; Probe: 5’ FAM-CATCAGAATCTGTAGGCCGT-3’ TAMRA]. In Q-RT-PCR analysis, the standard plasmid harboring the F3L gene was used to establish a standard curve, enabling the absolute quantification of viral copy numbers. The Q-RT-PCR were run on the ABI 7500 System (ThermoFisher, Waltham, MA, USA).

### Statistical analysis

Data were analyzed using GraphPad Prism software (San Diego, CA, USA). One-way analysis of variance (ANOVA) was used to determine statistically significant differences between groups. *P* < 0.05 was considered statistically significant. All assays were performed in triplicate and are representative of at least three independent experiments.

## Results

### Body weight changes and survival rate of dormice following MPXV infection

We evaluated the pathogenicity of MPXV in dormice. Following intranasal inoculation with MPXV, the animals exhibited a significant downward trend in body weight, with a peak reduction of 14.1% compared to their initial weight on 10 dpi (Figure 1A). The peak mortality period occurred between days 6 and 8 post-infection, resulting in a final mortality rate of 70% (Figure 1B). In contrast, the body weight of dormice in the control group increased gradually, and no deaths were observed throughout the entire experimental period (Figure 1).

**Figure 1.**
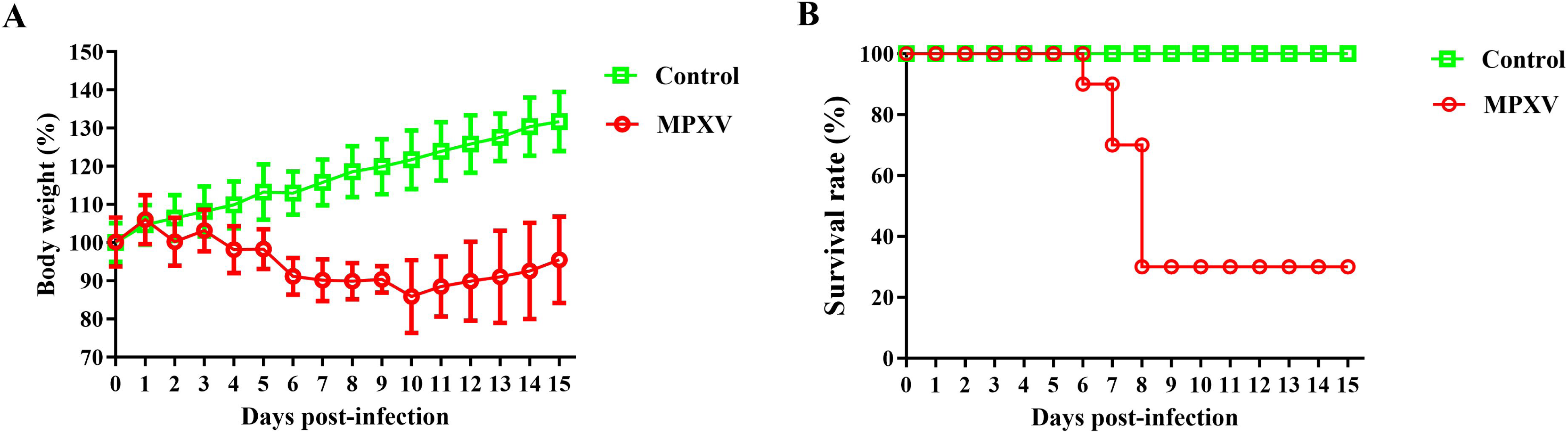
Body weight and survival rate in dormice infected with monkeypox virus. Ten healthy male dormice, aged 10 to 12 months, were selected and intranasally inoculated with the monkeypox virus (MPXV) to establish the virus-infected group. A separate group of ten animals was administered an equivalent volume of DMEM as a control. Over a 15-day period, daily monitoring was performed to assess body weight changes (A) and survival rates (B) in both groups.

### Tissue distribution of MPXV in dormice

We conducted a comprehensive investigation into the diffusion and replication kinetics of MPXV within the tissues of dormice. Our findings revealed that, following intranasal infection with MPXV, the virus rapidly disseminated to multiple organs throughout the animal’s body. Nucleic acid testing confirmed the presence of viral nucleic acids in all organs of the infected dormice, suggesting that the virus may disseminate systemically via the bloodstream to all organs (Figure 2A). Subsequently, infectious viruses were robustly detected in the nasal turbinates, trachea, lungs, and liver, with peak titers observed on the 7 dpi (Figure 2B). In contrast, infectious viruses were sporadically detected in the kidneys, spleen, and intestines, while no infectious virus was detected in the heart, brain, or testes throughout the experiment. These results suggest that the nasal turbinates, trachea, lungs, and liver may serve as the primary target organs for MPXV infection in dormice.

**Figure 2.**
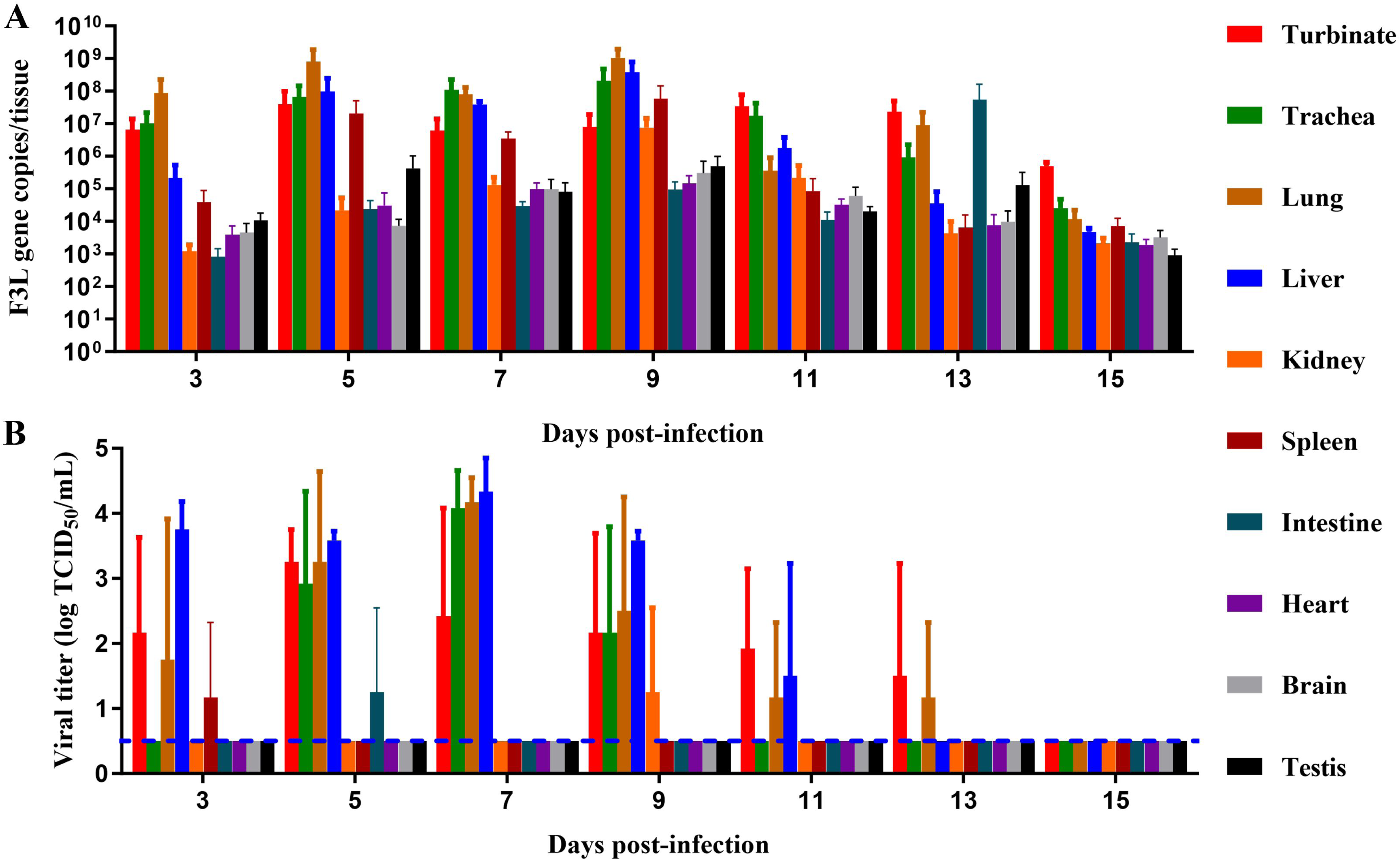
Tissue distribution of MPXV in dormice. Twenty-one male dormice, aged 10-12 months, were intranasally inoculated with MPXV. Three animals per time point were humanely euthanized on days 3, 5, 7, 9, 11, 13, and 15 post-infection, respectively. The nasal turbinates, trachea, lungs, liver, kidneys, spleen, intestine, heart, brain, and testes were collected and homogenized in PBS. The samples were then centrifuged, and the supernatants were collected for further analysis. Subsequently, the viral nucleic acid quantity (A) and viral titer (B) were quantified for each sample. Each color-coded column represents a distinct organ. Data are presented as mean ± SD. The dashed lines indicate the lower limit of detection.

### MPXV infection-induced pathological tissue damage in dormice

We further investigated the histopathological changes in dormice following MPXV infection (Figure 3). In the infection group, only the nasal turbinate (Figure 3A), trachea (Figure 3B), and spleen (Figure 3F) exhibited no significant pathological damage. Other organs, including the lungs, liver, kidneys, intestine, heart, brain, and testis, displayed varying degrees of pathological injury. In contrast, no obvious pathological damage was observed in the tissues of the negative control group.

**Figure 3.**
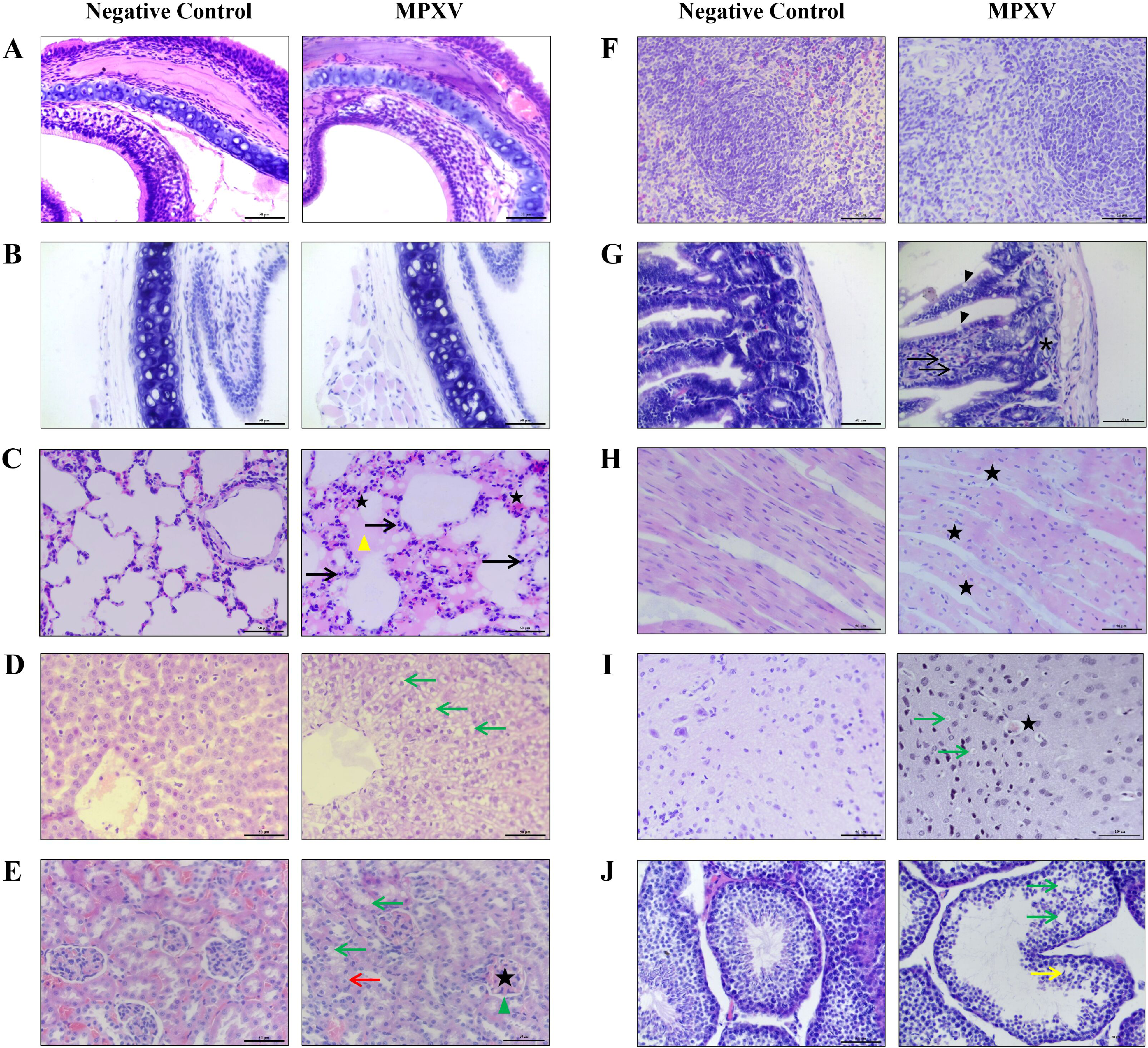
Histopathological analysis of multi-organ damage following infection with monkeypox virus (MPXV) Male dormice aged 10 to 12 months were selected and intranasally inoculated with either monkeypox virus (MPXV) or DMEM, serving as the virus-infected group and the negative control group, respectively. On the 7 dpi, multiple organ tissues were collected from both groups of dormice for analysis, including (A) nasal turbinate, (B) trachea, (C) lungs, (D) liver, (E) kidneys, (F) spleen, (G) small intestine, (H) heart, (I) brain, and (J) testes. The collected samples were fixed in formalin, embedded in paraffin, and subsequently subjected to pathological examination via hematoxylin-eosin (H&E) staining. The pentagram symbol denotes vascular or capillary hyperemia, encompassing glomerular vascular hyperemia and dilation. Black arrows denote the infiltration of inflammatory cells. Yellow triangles represent the presence of serous exudates. Green arrows indicate cytoplasmic vacuolization or presence of vacuoles between cells. Red arrows indicate cell granular degeneration. The green triangle indicates narrowing of the Bowman’s capsule cavity. Black triangles indicate an increased number of goblet cells. Asterisks indicate a decrease in the small intestinal glands of the small intestine. The yellow arrow indicates the reduction of spermatids in the seminiferous tubules. The scale represents 50 μm or 100 μm.

(Figure 3C) Lung: In the control group, the lung tissue structure was normal and intact. In contrast, in the infected group, the lung interstitium exhibited a significant increase in width, and the alveolar walls demonstrated varying degrees of thickening. Pulmonary interstitial capillaries exhibited congestion (adjacent to the pentagram), accompanied by inflammatory cell infiltration (indicated by black arrows) and serous exudates within the alveolar spaces (marked by yellow triangles).

(Figure 3D) Liver: In the control group, the structure of hepatocytes, hepatic cords, and hepatic sinusoids was relatively well-preserved. In contrast, the infected group exhibited significant vacuolar degeneration of hepatocytes(highlighted by green arrow), hepatocyte swelling, and a marked narrowing of hepatic sinusoids.

(Figure 3E) Kidney: In the control group, the renal corpuscles exhibited normal morphology, and no histopathological alterations were observed in the renal tubular epithelial cells. In comparison to the control group, the infected group demonstrated mild hyperemia and swelling of the renal glomeruli (indicated by pentacle), narrowing of the Bowman’s capsule cavity (indicated by green triangle), granular degeneration of renal tubular epithelial cells(highlighted by red arrow), and cytoplasmic vacuolization (highlighted by green arrow).

(Figure 3G) Intestine: In the control group, the intestinal villi structure was intact, and the mucosa, submucosa, and muscularis layers exhibited a well-defined stratified structure. In comparison to the control group, the infection group demonstrated a significant increase in the number of goblet cells (indicated by black triangles) within the intestinal mucosa, accompanied by slight thickening of the muscularis layer. In addition, lymphocytic infiltration within the lamina propria (as indicated by the black arrow) and a reduction in the number of small intestinal glands (as marked by the asterisk) were observed.

(Figure 3H) Heart: In the control group, the myocardium exhibited a uniformly consistent color, with myocardial fibers arranged in an orderly manner and a distinct, wide interstitial space between myocytes and muscle fibers. In contrast, the hearts of the infection group demonstrated markedly uneven myocardial coloration, disorganized arrangement of myocardial fibers, and significant myocyte swelling. Capillaries exhibit signs of congestion (adjacent to the pentagram).

(Figure 3I) Brain: In the control group, the brain exhibited a normal structure of the cerebral cortex and healthy neurons. In contrast, in the infection group, some capillaries within the cerebral cortex were observed to be congested (indicated by pentagrams). Additionally, neuronal degeneration was evident, characterized by pyknosis or loss of nuclei and cytoplasmic vacuolization (highlighted by green arrows).

(Figure 3J) Testis: In the control group, spermatogenic cells at all levels within the seminiferous epithelium of the seminiferous tubules were arranged in a clear and orderly manner. In the infected group, the number of spermatogenic cells in the seminiferous epithelium was decreased, and intercellular vacuoles (indicated by green arrows) were observed within the seminiferous epithelium. Compared with the control group, the structure of the seminiferous epithelium was disorganized, and severe exfoliation of spermatogenic cells was evident. A significant reduction was observed in the number of sperm within the seminiferous tubules (indicated by the yellow arrow).

### Direct contact and airborne transmissibility of MPXV in dormice

We performed a comprehensive evaluation of the transmissibility of MPXV among dormice through both direct contact and airborne transmission routes. We successfully detected viral nucleic acid in the respiratory tissues of all animals within the direct contact transmission group (Figure 4A), thereby demonstrating that MPXV can be transmitted between dormice through direct contact at the molecular level. On day 13 post-exposure, infectious viruses were detected in one-sixth of the recipient dormice (1/6) (Figure 4B). Additionally, on day 16 post-exposure, infectious viruses were detected in one-third of the recipient dormice (2/6), and seroconversion was observed in two-thirds of the animals (4/6; Supplementary Figure 1A). These findings suggest that MPXV possesses the capability for direct contact transmission among dormice. Moreover, infectious viruses from the recipient dormice were detected solely in the nasal turbinates, indicating that infection likely occurred via the nasal cavity.

**Figure 4.**
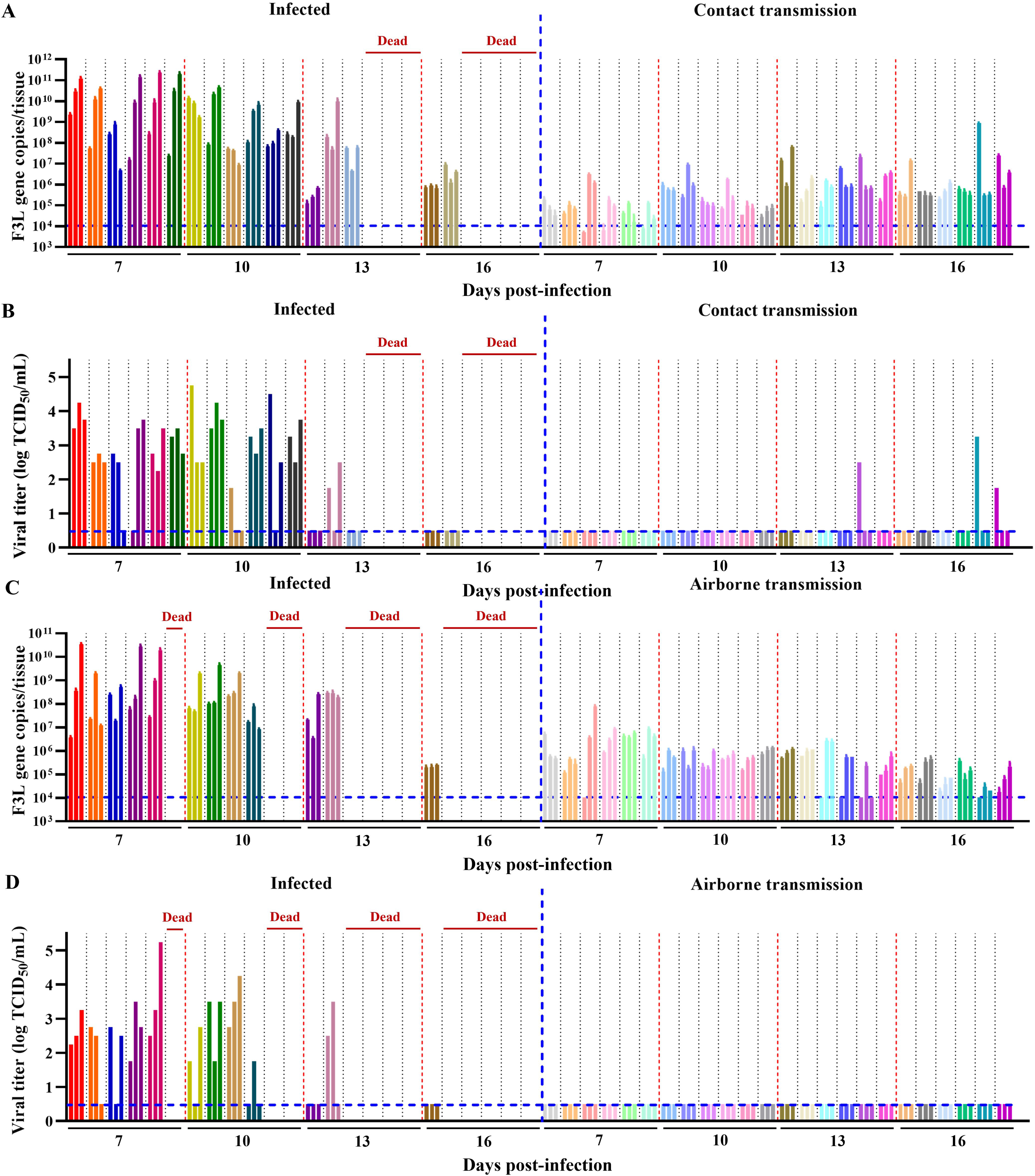
Direct contact and airborne transmissibility of MPXV in dormice. To assess the direct contact transmissibility, infected donor dormice were co-housed with direct contact recipient dormice, allowing for the direct physical contact. For the evaluation of airborne transmissibility, donor animals were placed in a wire-frame cage adjacent to airborne exposure recipient dormice. The distance between the infected donor dormice and the airborne-exposed dormice was 2 cm, which prevented direct contact while allowing indirect interaction via respiratory transmission. On days 7, 10, 13, and 16 post-infection or exposure, six donor or recipient dormice in each group were humanely euthanized. Thereafter, the nasal turbinates, trachea, and lungs were collected from these animals for quantification of viral nucleic acid copies and viral titers in the respective organs. The insets in the figure respectively illustrate the viral nucleic acid concentrations (A) and viral titers (B) in the respiratory tissues of dormice from the direct contact transmission experiment, as well as the viral nucleic acid concentrations (C) and viral titers (D) in the respiratory tissues of dormice from the airborne transmission experiment. Each set of three column bars in the same color represents the three respiratory tissues (from left to right: nasal turbinate, trachea, and lungs) of an individual dormouse. The dashed lines indicate the lower limit of detection.

We also successfully detected viral nucleic acid in the respiratory tissues of all animals within the airborne transmission group (Figure 4C), indicating that viral molecules can be transmitted via the airborne route. However, no infectious virus was detected in the airborne transmission group during the entire experimental observation period (Figure 4D). On day 16 post-exposure, one-third of the dormice became seropositive (2/6; Supplementary Figure 1B). These results suggest that although MPXV molecules are capable of airborne transmission, they are not sufficient to induce substantial infection. Taken together, MPXV exhibits limited airborne transmissibility among dormice.

### Concentrations and particle size distribution of viral aerosols exhaled by MPXV-infected dormice

In the airborne transmission experiment, we successfully detected viral nucleic acid in all airborne exposure recipient dormice, demonstrating that infected dormice can release viral aerosols. Furthermore, the viral particles present in exhaled aerosols are capable of transmitting between dormice, and inducing seroconversion in one-third of exposed dormice. Therefore, we further analyzed the viral shedding patterns in exhaled viral aerosols of MPXV-infected dormice. A six-stage Anderson impactor was utilized to collect the viral aerosols exhaled by the infected dormice on alternate day from 2 to 16 dpi. The total concentration of viral particles in the exhaled viral aerosols from dormice exhibited an increasing trend, reaching a peak at 12 dpi (3.78±1.01×10^6^ viral copies per dormice per hour’s breath), followed by a gradual decrease by 16 dpi (Figure 5A).

**Figure 5.**
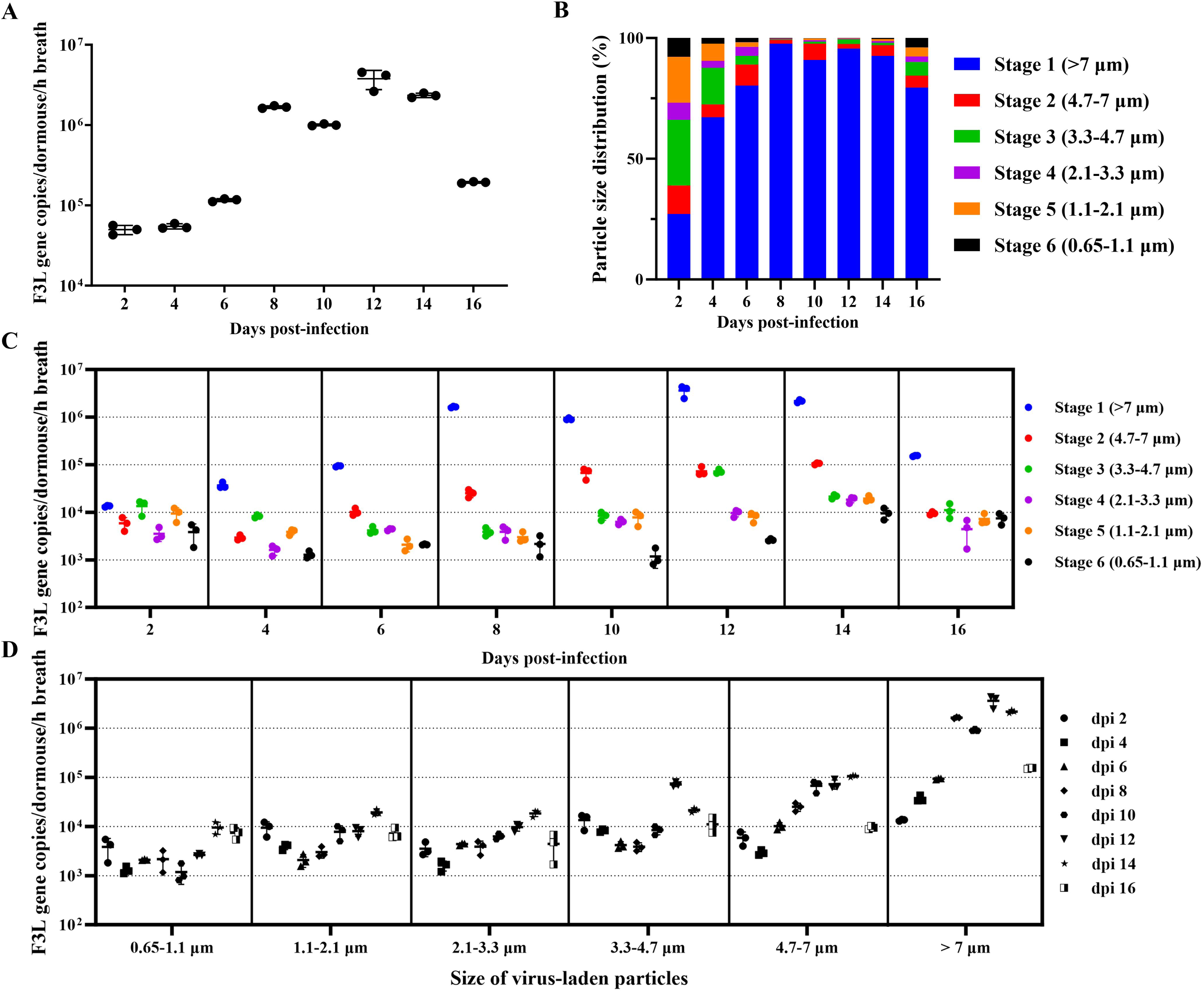
Concentrations and particle size distribution of virus-laden particles in the exhaled aerosols from MPXV-infected dormice. From 2 to 16 days post-infection, a six-stage Anderson impactor was employed to collect the virus-laden aerosols exhaled by the dormice on alternate day. (A) Total concentrations of virus-laden aerosols exhaled by MPXV-infected dormice. (B) Size distribution of virus-laden particles in the exhaled viral aerosols. (C) Quantity and size distribution of virus-laden particles exhaled by MPXV-infected dormice, categorized according to particle size range. (D) Quantity and size distribution of virus-laden particles exhaled by MPXV-infected dormice, categorized according to post-infection time points. Data in (A), (C), and (D) were overlaid with mean ± SD.

We further investigated the aerodynamic diameter distribution of virus-laden particles in viral aerosols exhaled by dormice. Our observations revealed that the exhaled aerosols contained a significantly higher abundance of larger-diameter particles compared to smaller ones (Figure 5B). During the peak phase of viral aerosol shedding (8-14 dpi), the majority of virus-laden particles in the viral aerosols exhaled by dormice were larger than 7 μm in diameter, accounting for over 90.9% of the total size distribution. Particles within the size range of 0.65 to 7 μm constituted only a minor fraction of the overall distribution.

Figure 5C and Figure 5D provide a more detailed analysis of the quantity and size distribution of virus-laden particles in the viral aerosols exhaled by dormice. The quantity of virus-laden particles with a size exceeding 7 μm remained consistently at approximately 10^6^ to 10^7^ copies per dormice per hour’s breath during the middle and late stages of infection (8 to 14 dpi). This was followed by the quantity of virus-laden particles within the size range of 4.7 to 7 μm, which was maintained at approximately 10^5^ copies per dormice per hour’s breath. The concentration of virus-laden particles with a diameter smaller than 4.7 μm remained consistently low throughout the infection period, rarely exceeding 10^4^ copies per dormouse per hour’s breath. As time progressed, the concentration of virus-laden particles in exhaled viral aerosols increased predominantly in coarse particles larger than 4.7 μm in diameter, while the quantity of fine particles ranging from 0.65 to 4.7 μm exhibited relatively minor fluctuations (Figure 5D).

### Median infective dose (ID_50_) of MPXV for intranasal infection in dormice

Finally, we conducted an in-depth investigation into the susceptibility of dormice to MPXV via respiratory tract infection by administering varying doses of MPXV, ranging from low to high, intranasally to groups of five dormice each. The maximum viral titer in the respiratory system (nasal turbinates, trachea and lungs) of each dormouse is presented in Figure 6A, while the infection rate for each group of dormice is summarized in Table 1. The infection rate of dormice and the viral titer in the respiratory system were observed to follow an infectious dose-dependent pattern. When the infectious dose was below 10^2.3^ TCID_50_, the infection rate in dormice remained at a relatively low level. Upon reaching a dose of 10^3.3^ TCID_50_, the infection rate increased significantly. At a dose of 10^4.3^ TCID_50_, the infection rate approached 100%. Probit analysis was performed to estimate the ID_50_ of MPXV infection in dormice (Figure 6B). The ID_50_ for MPXV infection in dormice was determined to be 10^2.773^ TCID_50_, with a 95% confidence interval ranging from 10^1.958^ to 10^3.827^ TCID_50_.

**Figure 6.**
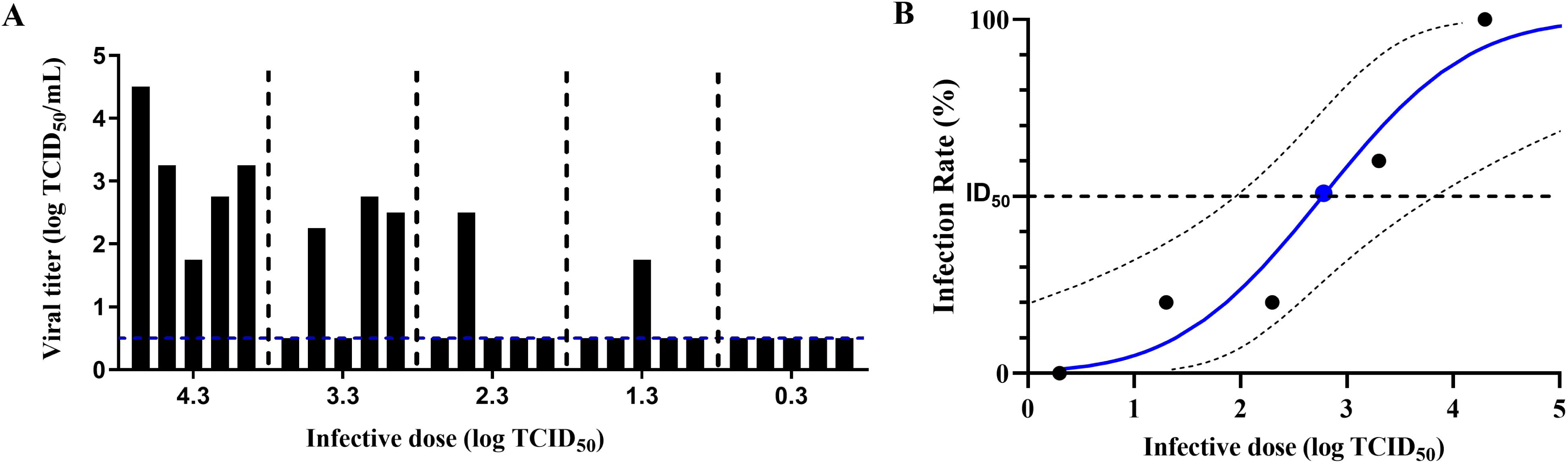
Median infective dose (ID_50_) of MPXV for nasal instillation infection in dormice. (A) Groups of five dormice were inoculated with the serially diluted MPXV. At 7 days post-infection (dpi), the dormice were euthanized, and their respiratory tissues, including the nasal turbinates, trachea, and lungs, were collected to determine the viral titers. Each bar represents the maximum viral titer detected in the respiratory tissues of each dormouse. Dashed line indicates the lower limit of virus detection. (B) Probit analysis was performed to determine the ID_50_ for instilled MPXV. Dashed lines show 95% confidence interval for the analysis.

**TABLE 1.** Infection rates of dormice inoculated with varying doses of MPXV.

| Infective dose<br>(lg TCID<sub>50</sub>) | Number of<br>infected dormice | Total dormice<br>population | Infection rate<br>(%) |
| --- | --- | --- | --- |
| 0.3 | 0 | 5 | 0 |
| 1.3 | 1 | 5 | 20 |
| 2.3 | 1 | 5 | 20 |
| 3.3 | 3 | 5 | 60 |
| 4.3 | 5 | 5 | 100 |

## Discussion

The monkeypox virus has traditionally been classified into two major genetic clades: the Central African clade (Congo Basin, CB) and the West African clade (WA). Research indicates that the pathogenicity of the CB clade is generally more pronounced compared to that of the WA clade(29). Schultz et al. investigated the effects of intranasal inoculation of the CB clade of monkeypox virus (MPXV-ZAI-79) and demonstrated that this virus is highly lethal to dormice, leading to substantial weight loss and severe clinical manifestations(27). The Monkeypox virus (MPXV-ZAI-79) was isolated in 1979 from a fatal human case in the Democratic Republic of the Congo. However, the ongoing outbreak is genetically closely associated with monkeypox viruses belonging to the hMPXV-1A lineage, which falls under the WA clade(30). Since May 2022, the WA clade has been identified as the primary contributor to the global monkeypox outbreak(20). Therefore, we utilized the WA clade monkeypox virus (hMPXV/China/CCVRI-01/2023) to develop an animal model and investigate the pathogenicity and transmissibility of the monkeypox virus. In contrast to prior studies, we observed a comparable tissue distribution of MPXV in dormice. Nevertheless, in our study, the WA clade monkeypox virus exhibited a lower mortality rate in dormice. Among the pathological findings, no inflammatory effects of the CB clade of MPXV were observed on the nasal mucosa or lymphatic necrosis in the spleen of dormice. These differing results further highlight the variations in pathogenicity of MPXV across its various clade.

The Monkeypox virus (MPXV) was originally endemic to Central and Western Africa. However, in recent years, it has exhibited a rapid global expansion facilitated by various transmission pathways, including animal-to-animal spread, cross-species infection, and human-to-human transmission. According to the U.S. Centers for Disease Control and Prevention (CDC), the monkeypox virus can be transmitted through various modes of human-to-human transmission. These include, but are not limited to, sexual contact, direct contact with affected areas such as rashes, ulcers, or scabs, contact with contaminated objects like clothing or bedding, and inhalation of respiratory droplets, aerosols, or oral secretions from an infected individual(31). Although sexual contact transmission is regarded as the primary driver of the current monkeypox outbreak, the precise role and contribution of other transmission routes in the dissemination of the epidemic warrant further investigation(32). Furthermore, considering the unique characteristics of the monkeypox virus as a potential bioterrorism agent, its efficiency and capacity to transmit among humans via respiratory secretions, including droplets, aerosols, or exhaled air, warrant significant attention(33). Therefore, it is of paramount urgency to investigate the transmission dynamics of the monkeypox virus and conduct a systematic assessment of the impact of various transmission routes on the epidemic spread, given the increasingly severe transmission of the monkeypox virus.

There is currently a notable absence of systematic research regarding the transmissibility of monkeypox viruses across different animal populations. To date, apart from studies involving black-tailed prairie dogs, no direct investigations into monkeypox virus transmission within susceptible animal populations have been documented in the literature. Hutson et al. conducted an experiment in which a black-tailed prairie dog infected with the WA strain (MPXV-USA-2003-044) via intranasal inoculation was co-housed with three other healthy black-tailed prairie dogs to simulate the contact transmission process of MPXV(25). The results demonstrated that the viable virus could be detected in oral secretions as early as 13 days post-exposure in three prairie dogs exposed via the contact route. Similarly, in our study, the process of contact transmission of the WA strain (hMpxV/China/CCVRI-01/2023) between dormice was simulated using co-housing conditions. At the corresponding time point (13 days post-infection), viable infectious virus was successfully isolated from the turbinates of the recipient dormice.

It is worth noting that, regardless of whether it is the black-tailed prairie dogs or the dormice, the infectious virus was initially detected in the upper respiratory tract of naïve recipient animals. In addition, none of the naïve recipients housed in the same cage exhibited any signs of biting or scratching behavior. Therefore, we posit that the transmission of the monkeypox virus is primarily facilitated through respiratory secretions, such as exhaled breath and droplets originating from the mouth and nose. These secretions serve as critical vectors in the dissemination of the monkeypox virus. Studies have demonstrated that black-tailed prairie dogs are capable of transmitting the monkeypox virus exclusively through the exchange of exhaled air and respiratory secretions(25).

Hutson et al. conducted an experiment in which six prairie dogs experimentally infected with the monkeypox virus were placed in cages positioned opposite six naive recipient prairie dogs, maintaining a distance of 3 inches between the cages. There was no direct physical contact between the donor and recipient animals(26). The six prairie dogs experimentally infected with the monkeypox virus comprised three prairie dogs infected with the WA strain (MPXV-USA-2003-044) and three prairie dogs infected with the CB strain (MPXV-RCG-2003-358). Notably, one prairie dog was infected with a high dose of the CB strain, while the other two were infected with a low dose. Following an extended observation period, it was determined that among the six naïve recipient prairie dogs, only one individual became infected. This recipient prairie dog was positioned directly opposite the donor animal inoculated with a high dose of the CB strain virus. The remaining five recipient prairie dogs, comprising three housed opposite the donor animals infected with the WA strain and two housed opposite the donor animals infected with a low dose of the CB strain of monkeypox virus, remained uninfected. This study demonstrates that, in comparison to direct contact transmission, the respiratory route is a less efficient mode of monkeypox virus transmission among black-tailed prairie dogs. In our study, a comparable methodology was utilized to simulate the respiratory transmission process of the monkeypox virus among dormice, and consistent outcomes were observed. MPXV exhibits limited airborne transmissibility among dormice. Although MPXV molecules are capable of airborne transmission, they are not sufficient to induce substantial infection.

The transmission characteristics of the monkeypox virus in dormice are comparable to those observed in humans. The transmission of the monkeypox virus among dormice is contingent upon direct and close contact between the animals. According to epidemiological surveillance data, the chain of human-to-human transmission of the monkeypox virus is significantly dependent on direct contact(34). Among unvaccinated close contacts, the secondary attack rate among family close contacts of the primary case ranged from 7.5% to 12.3%, whereas the secondary attack rate among non-family close contacts was significantly lower at 3.3% (35, 36). At the same time, the extension of the transmission chain is evidently constrained, and tertiary transmission events (contact infections resulting from secondary infection cases) as well as quaternary transmission events (contact infections resulting from tertiary infection cases) are only sporadically documented in existing epidemiological reports (37, 38). The airborne transmission capacity of the monkeypox virus among dormice appears to be restricted. Likewise, person-to-person airborne transmission of the monkeypox virus remains unconfirmed. To date, only limited and indirect evidence exists to suggest the potential for airborne transmission(39, 40). However, this possibility cannot be entirely dismissed at present(41, 42). Additional cases of monkeypox virus infection or further research into its transmissibility are required to confirm this hypothesis. Further research utilizing additional animal models is essential for a comprehensive understanding of the transmissibility dynamics of the monkeypox virus.

There are potentially two reasons for the limited airborne transmission capability of the monkeypox virus among dormice. On the one hand, the 50% infectious dose of the monkeypox virus through respiratory tract infection is relatively high compared with other respiratory pathogens. We determined the 50% infection dose of monkeypox virus in dormice infected by nasal drops and found that it was around 10^2.7^ TCID_50_. The 50% infection dose of pathogenic microorganisms transmitted via the respiratory route is typically around 10^1^ TCID_50_, as observed with the influenza virus and SARS-CoV-2(43). In contrast, the 50% infectious dose of the monkeypox virus via respiratory infection is relatively high, which provides an initial explanation for the limited airborne transmission capacity of the monkeypox virus within dormice populations.

In addition to the susceptibility of host animals to pathogenic microorganisms, the particle size of airborne pathogenic microorganisms also plays a critical role in determining their capacity for airborne transmission(44). The propagation distance of droplets larger than 4.7 μm is relatively limited. In contrast, smaller particles measuring less than 4.7 μm remain suspended in the air for extended periods, disperse over wider areas, and are more readily inhaled deep into the respiratory tracts of humans or animals. Zaucha et al. (45) and Barnewall et al. (46) independently aerosolized the CB strain (Zaire V79-I-005) into aerosols with median particle sizes of 1.2 micrometers and 1.08–1.15 micrometers (mass median aerodynamic diameter), respectively, thereby establishing an inhalation monkeypox virus infection model in cynomolgus monkeys. This small size range of monkeypox virus aerosol particles can successfully reach the alveoli of cynomolgus monkeys and cause infection in the animals. In the present study, the median size of viral particles in the exhaled aerosol of dormice infected with monkeypox virus was greater than 7 μm. These coarse particles are likely to travel only a limited distance through the air and may not readily pass through the nasal cavity of dormice into the lungs, instead predominantly settling in the upper respiratory tract of the animals. One of the reasons for the relatively low efficiency of respiratory transmission of the monkeypox virus among dormice is the larger particle size of the virus-laden aerosols exhaled by infected dormice.

Our study demonstrates that African dormice are susceptible to the monkeypox virus and highlights the potential for transmission of the virus to other uninfected animals through contact within dormouse populations. African rodents serve as critical components in the natural transmission cycle of the monkeypox virus. Acting as natural hosts, they play a pivotal role in maintaining and disseminating the virus within ecosystems(47).In certain regions of Africa, human infection with the monkeypox virus through contact with these virus-carrying rodents has been reported frequently. The ecological behaviors of these rodents facilitate the cross-regional and cross-species transmission of the virus(48). Wild African dormice populations serve as a critical component in the natural transmission cycle of the monkeypox virus. On the one hand, dormice serve as natural hosts that are capable of replicating the monkeypox virus following infection. These animals can subsequently carry and shed the virus into the external environment over extended periods, thus sustaining the presence of the monkeypox virus in natural ecosystems. On the other hand, infected dormice serve as critical vectors for the transmission of the monkeypox virus.

Following infection, dormice are capable of disseminating the virus to other animals within the same region during their routine activities, including various rodent species and non-human primates. This facilitates the circulation and propagation of the monkeypox virus throughout the ecosystem, consequently elevating the likelihood of zoonotic spillover events leading to human infections. In addition, dormice kept as pets may serve as a significant link in the transmission chain of the monkeypox virus during international trade, thereby potentially posing a considerable threat to public health(49).

In summary, the African dormouse serves as a suitable animal model for assessing monkeypox virus infection, pathogenicity, and transmission dynamics. Dormice infected with the monkeypox virus display distinct pathogenic features, and the virus can be transmitted between populations via direct contact. Although the virus exhibits relatively limited aerosol transmission capability, experimental evidence has confirmed that infected dormice can exhale substantial amounts of viral aerosols, thereby posing a potential public health risk. At the same time, intensified surveillance of wild dormice populations is essential given their possible role in monkeypox virus transmission chains. Finally, it is recommended to develop and implement appropriate regulatory frameworks in order to effectively disrupt any potential chain of animal-to-human transmission. These findings offer critical evidence for enhancing and reinforcing the prevention and control strategies against monkeypox.

## Acknowledgement

We would like to express our gratitude to the Eighth Affiliated Hospital of Guangzhou Medical University for providing clinical samples of monkeypox.

## Authorship contribution statement

**Zhaoliang Chen:** Writing – original draft, Methodology, Formal analysis, Conceptualization. **Lei Zhang:** Methodology, Validation, Investigation. **Linzhi Li:** Methodology, Validation, Investigation. **Mingjie Shao:** Methodology, Validation, Investigation. **Zhongzeng Zhao:** Investigation, Formal analysis. **Shang Chao:** Methodology, Investigation. **Zirui Liu:** Methodology, Investigation. **Juxiang Liu:** Methodology, Investigation. **Yan Liu:** Conceptualization, Writing – review & editing, Project administration. **Xiao Li:** Conceptualization, Writing – review & editing, Project administration. **Zhendong Guo:** Writing – review & editing, Writing – original draft, Supervision, Project administration, Validation, Conceptualization.

## Data availability statement

The data will be made available upon request.

